# Predicting compound-protein interaction using hierarchical graph convolutional networks

**DOI:** 10.1101/2021.10.04.463093

**Authors:** Danh Bui-Thi, Emmanuel Rivière, Pieter Meysman, Kris Laukens

**Affiliations:** Adrem Data Lab, University of Antwerp, Antwerp, Belgium

## Abstract

**Motivation:** Convolutional neural networks have enabled unprecedented breakthroughs in a variety of computer vision tasks. They have also drawn much attention from other domains, including drug discovery and drug development. In this study, we develop a computational method based on convolutional neural networks to tackle a fundamental question in drug discovery and development, i.e. the prediction of compound-protein interactions based on compound structure and protein sequence. We propose a hierarchical graph convolutional network (HGCN) to encode small molecules. The HGCN aggregates a molecule embedding from substructure embeddings, which are synthesized from atom embeddings. As small molecules usually share substructures, computing a molecule embedding from those common substructures allows us to learn better generic models. We then combined the HGCN with a one-dimensional convolutional network to construct a complete model for predicting compound-protein interactions. Furthermore we apply an explanation technique, Grad-CAM, to visualize the contribution of each amino acid into the prediction.

**Results:** Experiments using different datasets show the improvement of our model compared to other GCN-based methods and a sequence based method, DeepDTA, in predicting compound-protein interactions. Each prediction made by the model is also explainable and can be used to identify critical residues mediating the interaction.

**Availability and implementation:** https://github.com/banhdzui/cpi_hgcn.git

## Introduction

The identification of compound-protein interactions (CPI) plays an important role in drug discovery and development. Typical applications include high-throughput screening of compound libraries for given protein targets to identify desirable compound-protein interactions or testing given compounds against possible off-target proteins to avoid undesired effects [1, 2]. Experimental identification of every possible compound and protein pair is unpractical, if not impossible, due to the enormity of the chemical and proteomic space. Therefore, computational methods to predict compound-protein interactions have received increasing attention. Especially the adaptation of deep learning models to structured data has opened a new paradigm for pharmaceutical research.

Given a compound-protein pair, CPI prediction methods aim to predict a binary value indicating whether the compound and the protein interact [3–6], a numeric value indicating their binding affinity [1, 7–12], or identify binding sites for a specific compound within the protein [13–16]. Existing CPI prediction methods are diverse in terms of feature engineering and machine learning models. They can be categorized into different classes. The first class consists of similarity-based models, which utilize drug-drug and target-target similarity matrices to infer possible interactions. This approach commonly applies nearest-neighbor and kernel-based classifiers as machine learning models. A non-exhaustive list of such methods includes [17–20]. While similarity measures vary greatly among methods, drug-drug similarity is generally based on chemical structure fingerprints, whereas target-target similarity typically depends on a sequence alignment score [8]. In the second class, several studies [21, 22] have analyzed binding affinity matrices for interaction prediction. Matrix factorization techniques have been used in this case to reveal latent features for drugs and proteins. A study from 2017 [8] presented SimBoost, combining both similarity matrices and binding affinity matrices to construct features for drugs, targets and drug-target pairs. These features were used as the input of a Gradient Boosting Regression Tree, creating a model to predict binding affinity. The third class consists of learning models that utilise pre-defined features and fingerprints to classify binders from non-binders. For example, Cheng et al. [23] trained support vector machine (SVM) models, using Molecular ACCess System (MACCS) keys and 1400 PROFEAT protein descriptors as input features. Likewise, Yu et al. [24] used PROFEAT protein descriptors in combination with chemical compound descriptors to train Random Forest and SVM models. Lastly, Wen et al [25] built a Restricted Boltzmann Machine with Extended Connectivity Fingerprints (ECFP) and protein sequence composition.

As a result of important theoretical and practical developments in recent years, several deep learning based methods have been devised to improve CPI identification. Deep learning models with their complex architecture are able to learn abstract features from raw data, which can lead to improved performance. Convolutional neural networks (CNN) have shown impressive performance in a variety of tasks, such as image recognition, text classification and audio processing [26]. Building on this success, several studies have investigated the use of a 3D-CNN model to predict CPI using 3D-structural data [27, 28]. Typically, the data is constructed by discretizing the protein-compound molecular structure into a 3D grid centered around the binding site. However, structural information regarding the molecular interaction between the protein and compound is not always available, limiting the scope of these methods. Recently, Zhao et al. [6] and Zong et al. [29] built association networks among drugs and targets, allowing them to learn features for drugs, targets or drug-target pairs using graph embedding learning algorithms such as graph convolutional networks (GCN) [30] or node2vec [31]. The drawback of these network-based methods is that they require retraining when new nodes are inserted and cannot predict associations for any unseen nodes.

Some of the most readily available data representations in CPI datasets are Simplified Molecular Input Line Entry System (SMILES) strings for compounds and amino acid sequences for proteins. Each SMILES string corresponds to a unique molecular graph that describes the structure of the compound. The nodes of these graphs represent atoms, while the edges represent the covalent bonds between atoms. Given the availability of large CPI data sets, several studies have investigated deep learning models that work directly with these data formats. In general, these models consist of three key components, each built from a set of distinct neural network layers. Two components are responsible for encoding compounds and proteins respectively, while the final component translates the output of the encoding layers into a CPI prediction. The predicting component is usually a fully-connected network, computing whether an interaction occurs or computing a binding affinity value. The protein is commonly encoded through sequence-based models. DeepDTA [11], DeepConv-DTI [9], GraphDTA [10], Tsubaki et al. [5], MT-DTI [12] and TransformerCPI [3] apply 1D-CNN layers to encode protein sequences, while DeepAffinity [1] and DeepCDA [7] combine 1D-CNN layers with recurrent neural network (RNN) or long short-term memory (LSTM) layers, respectively. The compound is encoded with sequence-based or graph-based models, depending on the input information. DeepDTA, DeepAffinity, DeepCDA, and MT-DTI developed sequence-based models which utilise SMILES strings directly, considering compounds as sequences of finite letters. In contrast, several studies have explored the molecular structure of compounds as input features. DeepConv-DTI [9] uses ECFP fingerprints combined with a fully-connected network while the studies [4, 5, 10] investigate in GCNs to learn structural features of compounds. Different GCN models have been implemented, including Graph Laplacian based GCN [32], Graph Attention Networks (GAT) [33] and Graph Isomorphism Networks (GIN) [34]. In the TransformerCPI method, a GCN was combined with Transformer Decoder, a sequence-based model, to generate an embedding for every atom of the compound. In 2019, Karimi et al. [1] compared performance of the two approaches for compound representation. They implemented a RNN working with SMILES string and the GCN proposed by the study [4]. Their result showed that the GCN does not outperform the RNN. We obtained the same result when we compared performance of a recent sequence-based model with some existing GCNs on *human, C. elegans* [35] and *bindingdb* [4] data. However, graph structures are more descriptive than SMILES strings, especially for model interpretation [1]. Therefore, developing a better graph-based model to represent compounds are still an active problem.

The deep learning methods for CPI prediction have shown to outperform non-deep learning methods on benchmark datasets [3, 5, 11]. In this study, we explore a novel deep learning approach to predict whether compounds and proteins interact, based on compound graphs and amino acid sequences. We focus more on building an efficient GCN to represent compound structure. Similar to existing deep learning methods, our model includes three components: a GCN to encode compounds, a 1D-CNN based model to encode proteins, and a fully connected network to predict CPI. We postulate that looking at the local parts of data objects independently can improve the performance of trained models, especially for small molecules, which usually share many substructures. From this rationale, we introduce a procedure to construct the hierarchical representation for compounds in which compounds are considered as graphs of substructures, while the substructures are considered as graphs of atoms. Based on this approach, we propose a hierarchical GCN to model compounds. The hierarchical GCN learns an embedding for atoms, substructures, and then entire graphs, respectively. For the 1D-CNN based model encoding proteins, we combine 1D-CNN layers with pooling layers as well as fully-connected layers. Validation experiments on four different CPI datasets show that the proposed method provides an improved performance compared to existing GCN-based and string-based models. In addition, we applied Grad-CAM [36] on our model to evaluate the contribution of specific regions within protein sequences to the prediction.

## Our method

### Data pre-processing

The input of our model consists of proteins, represented as amino acid sequences, and compounds, represented as graphs. Amino acid sequences are projected into sequences of integers in which each integer represents a unique amino acid. Compound graphs are generated from SMILES strings using RDKit [37]. Compound graphs are then broken into subgraphs to account for aromatic links and atomic bonds. Our compound breaking procedure can be summarized as follows: First, the compound is split into two parts, one consisting of aromatic links and the other of non-aromatic links. This separation creates new graphs that can have more than one connected component in which each connected component corresponds to one specific substructure. Then, the connected component finding algorithm is used to detect the substructures. Finally, virtual edges are added between the substructures with which they share at least one atom in the original graph. This procedure generates a hierarchical representation of compounds. Figure 1 is an example of constructing the hierarchical representation for the compound with formula *C*_25_*H*_25_*ClN*_6_*O*. The resulting representation is a reduced graph including eight nodes and seven virtual edges: two nodes are Benzenes, one is Quinazoline and the other nodes are non-aromatic substructures. These nodes are also graphs, of which the nodes are atoms and the edges are connections between the atoms. For compounds consisting of only aromatic or non-aromatic bonds, the hierarchical representation contains a single node corresponding to the entire original graph.

**Fig 1.**
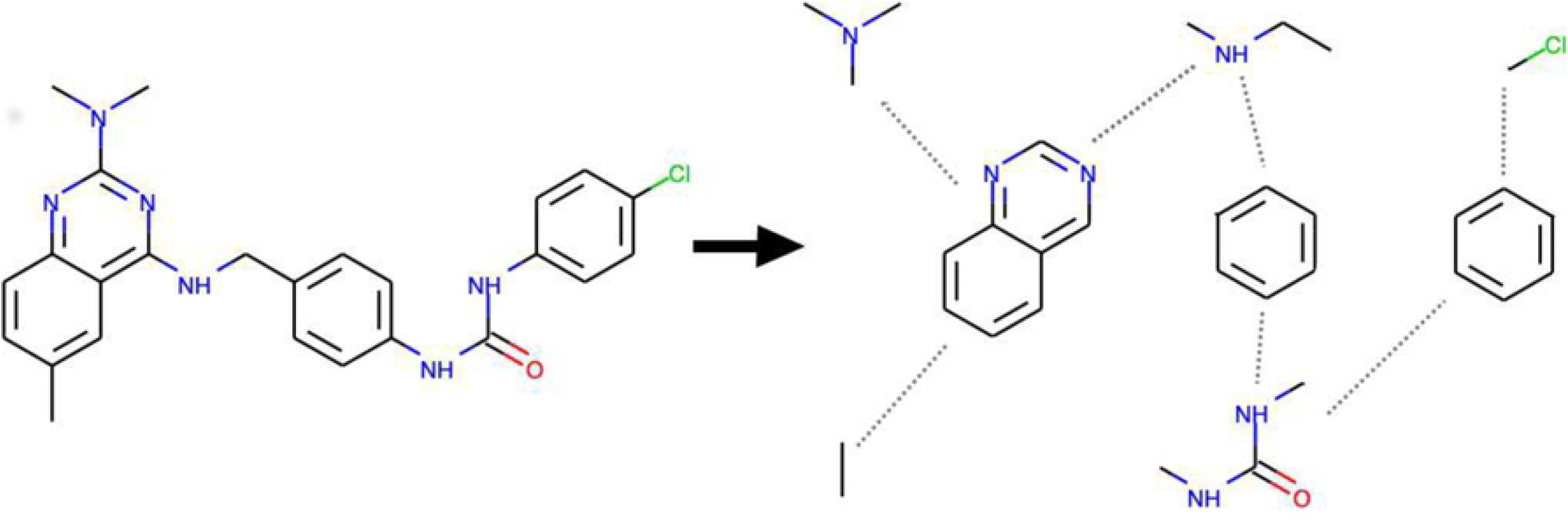
An example of constructing the hierarchical representation of *C*_25_*H*_25_*ClN*_6_*O* based on aromatic bonds and connected components.

### Compound representation

In this section, we present in detail the hierarchical graph convolutional network (HGCN) to encode compounds. In general, a graph convolutional network (GCN) is a neural network architecture defined according to graph structure 𝒢 = (𝒱, *ε*). Nodes *v* ∈ 𝒱 take unique values from 1,.., |𝒱| and edges in *ε* are pairs (*u, v*) ∈ 𝒱 × 𝒱. Graphs may contain node labels *l*_*v*_ ∈ {1,.., ℒ_𝒱_} for each node and edge labels *l*_(*u,v*)_ ∈ {1,.., ℒ_*ε*_} for each edge. Given initial node features *x*_*v*_ and edge features *e*_*uv*_, the aim of the GCN is to learn a state embedding *h*_*v*_ for every node *v* in 𝒢. The node embedding can then be used to synthesize the graph embedding of 𝒢. The forward propagation of GCNs includes two phases: message passing and readout [38]. The message passing phase computes node embedding through an iterative procedure which is defined in terms of message functions *M*_*t*_ and node update functions *U*_*t*_, as follows:

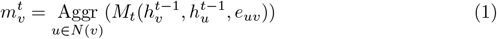

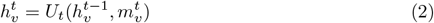

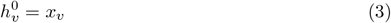

Here, Aggr denotes the aggregation function which can be a sum, mean or max function; *N* (*v*) are the neighbors of node *v* in graph 𝒢. As the functions *M*_*t*_ and *U*_*t*_ are typically shared over all locations in the graph, the neural network architecture is referred to as a convolutional neural network. At the readout phase, a function *R* is applied to compute the state embedding for the entire graph based on the node embedding at the last iteration *T*.

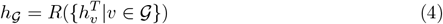

Existing GCN models vary in how these *M*_*t*_, *U*_*t*_, and *R* functions are defined.

Within the proposed HGCN, we compute the graph-level embedding based on the subgraph-level embedding which in turn is aggregated from the node-level embedding. The messages are passed around atoms within the subgraph scope using message functions 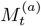 and update functions 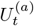. The communication among subgraphs is not considered in this step. After *T*^(*a*)^ iterations, the initial state embedding for every subgraph is assigned using a readout function *R*^(*sg*)^. Another message passing phase is running at the subgraph-level, in which messages are passed around through virtual edges with message functions 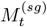 and update functions 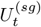. Finally, a readout function *R*^(𝒢)^ is exerted to aggregate the state embedding for the entire graph 𝒢. The node embedding is computed as follows:

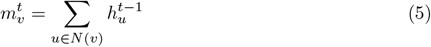

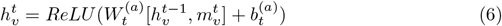

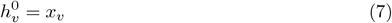

Here [*a, b*] denotes the concatenation of two vectors *a* and *b* in their last dimension; *x*_*v*_ are one-hot vectors corresponding to atom labels. The one-hot vectors can be replaced by pre-defined features of atoms. The embedding of every node *v* is concatenated with the signals from its neighbors and then the resultant vector is passed a fully-connected (FC) layer to obtain the update embedding of *v*. We choose concatenating over adding directly the neighbors’ signals to the embedding of *v* as they have different meanings and these meanings are learnable through learning the weights of the FC layer. At subgraph-level, we use the same message passing scheme at node-level but the initial state embedding for subgraphs *c*_*i*_ are aggregated from their atoms.

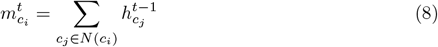

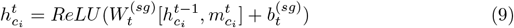

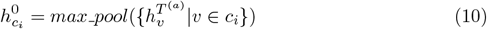

where *N* (*c*_*i*_) are the neighboring subgraphs of *c*_*i*_ according to the virtual edges; 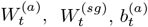 and 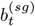 are learned parameters. After *T*^*sg*^ time steps, we compute graph embedding of 𝒢 according to:

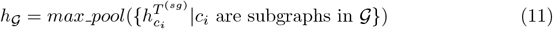

Within this study, we run the message passing phase at atom level with *T*^(*a*)^ = 2 time steps and at subgraph level with *T*^(*sg*)^ = 1 time steps. Each time step is associated with a neural network layer. Figure 2 illustrates the forward propagation of a HGCN for a simple graph which consists of two aromatic components, {*A, B, C*}, {*D, E, F*} and two non-aromatic components, {*A, G*}, {*B, E*}. *k* = 2 convolution layers at node-level continuously update embedding for every node based on the embeddings of their neighbors within subgraphs using the equations 5 and 6. Note that node *A* in the component {*A, B, C*} and node *A* in the component {*A, G*} are two different nodes that have the same label. As a result, their embedding are also different after the message passing phase, although they come from the same node in the original graph. When the *k*-th convolution layer completes, nodes are gathered to compute the embedding for subgraphs according to equation 10. For example, the embedding of the component {*A, G*} are computed from the embeddings of two nodes *A* and *G*. The convolution layer at subgraph-level takes into account the interactions between these components to update their embedding and finally generates the embedding for the entire graph.

**Fig 2.**
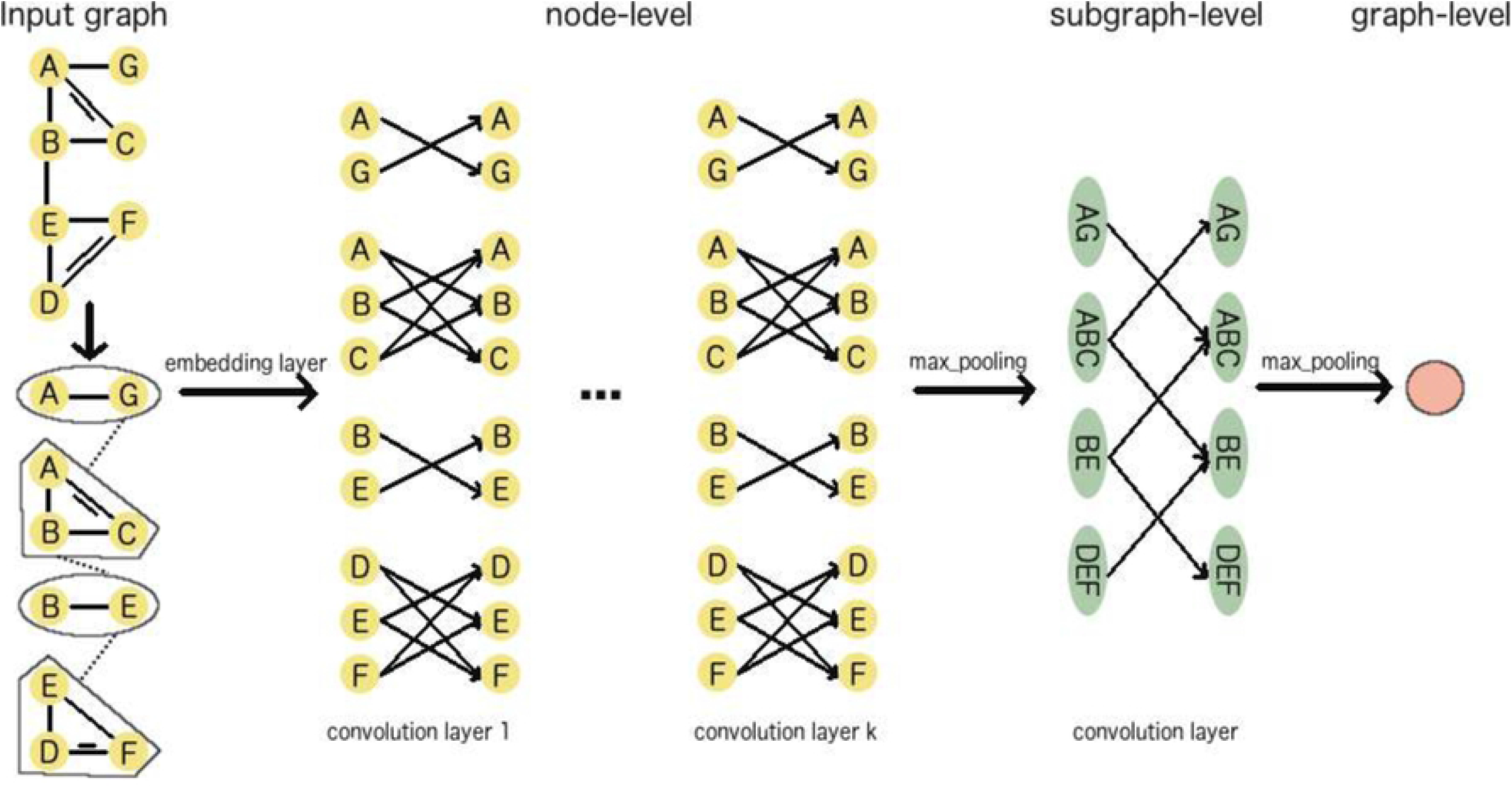
An illustration of the forward propagation of a HGCN which consists of *T*^(*a*)^ = *k* convolution layers at node-level and *T*^(*sg*)^ = 1 at subgraph-layer.

### Protein representation

Figure 3 demonstrates the model we built to encode protein sequences. We used a 1D-CNN based model, including three 1D-CNN layers of which two are followed by max pooling layers, and two fully-connected (FC) layers at the end. We fix the size of the kernels in the first 1D-CNN layer and the last max pooling layer to 3 and 12 respectively. For the other 1D-CNN and max pooling layers, this hyper-parameter can be customized by users. First, protein sequences are transformed into sequences of numerical vectors by an embedding layer, where every vector corresponds to an amino acid. These sequences then pass through the 1D-CNN and max pooling layers, downsampling into shorter sequences. When the last max pooling layer is finished, the resulting sequences are flattened into large vectors. The two FC layers at the end compress these vectors into smaller vectors that are the final representation of the protein.

**Fig 3.**
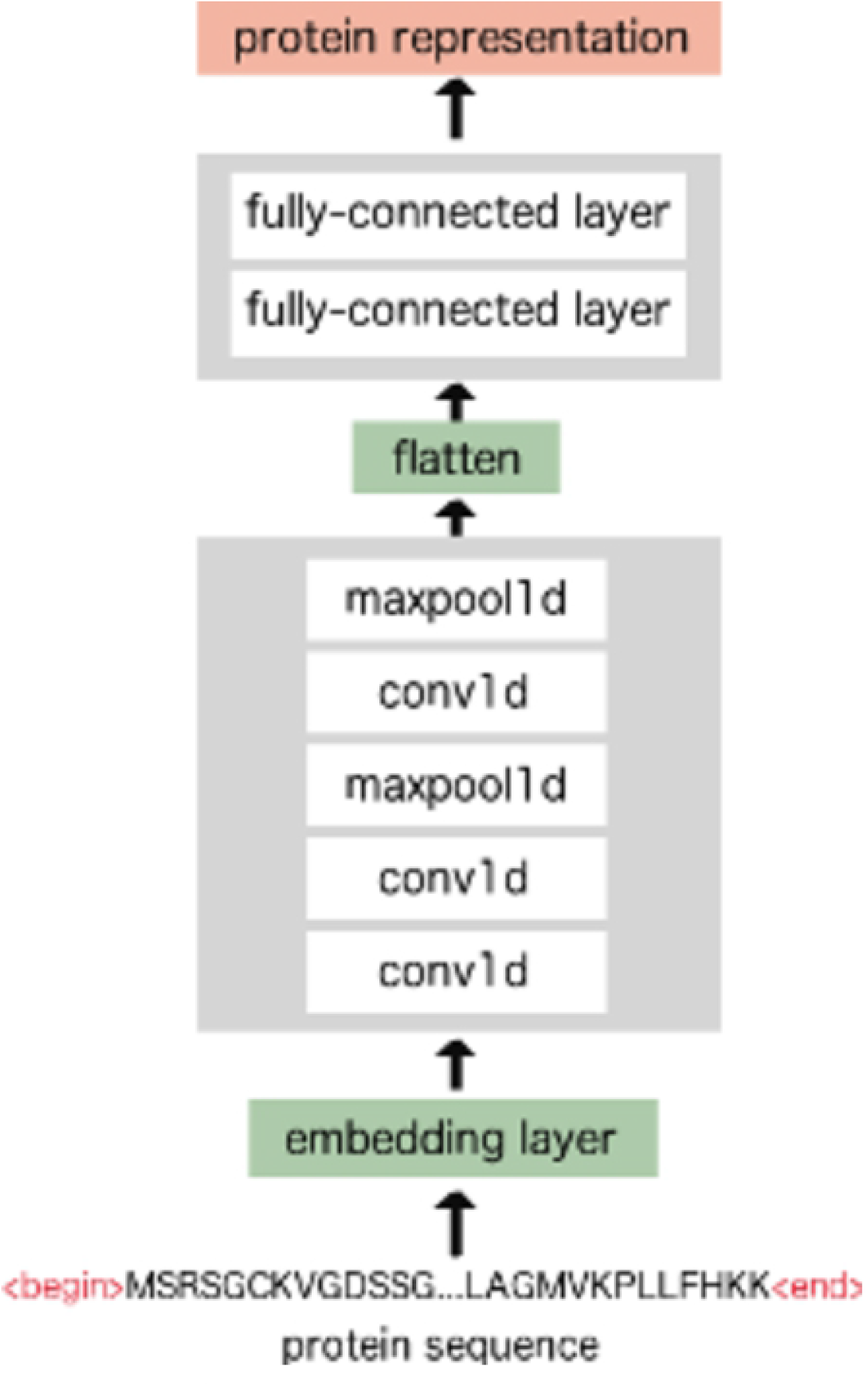
An overview of the 1D-CNN model that encodes protein sequences.

As protein sequences have varying length, we chose to fix the maximum length of input sequences using the length of the longest protein in the training data. The sequences that are longer than this maximum length are truncated, whereas shorter sequences are 0-padded. Additionally, we added two more special symbols into the input sequences in order to mark their begin and end position. As a result, the vocabulary for the embedding layer consists of 24 words in which 22 words correspond to amino acids and 2 words correspond to the begin/end symbols. The 22 amino acids include the 20 of the standard genetic code and an additional 2 that can be incorporated by special translation mechanisms, Selenocysteine (U) and Pyrrolysine (O).

### CPI prediction model

To predict a CPI, the compound embedding and the protein embedding is concatenated into a single vector. These vectors represent compound-protein pairs and are used as input features for a four layer fully-connected neural network (FCNN). The outcome of this network is a vector *z* ∈ ℛ^2^. A softmax layer is applied on top of the output vectors *z* to generate the CPI probability according to:

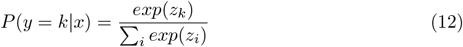

Here, *k* ∈ {0, 1} denotes class label, *x* is the set of compound-protein pair features and *z*_*k*_ is the value at the *k*-th dimension of vector *z. P* (*y* = *k*|*x*) is the probability for the *k*-th class (i.e. interact or not) given a CPI *x*. The compound embedding and protein embedding are generated by the HGCN and 1D-CNN based model described in section and, respectively. To reduce over-fitting, we applied dropout *p* = 0.2 for the two last layers of the 1D-CNN model. Figure 4 presents an overview of our CPI prediction model. In this study, all models are trained with learning rate *alpha* = 0.0001, dropout *p* and Adam [39] is used as the optimization algorithm in the training process. We implemented our model using the Pytorch and Pytorch Geometric library.

**Fig 4.**
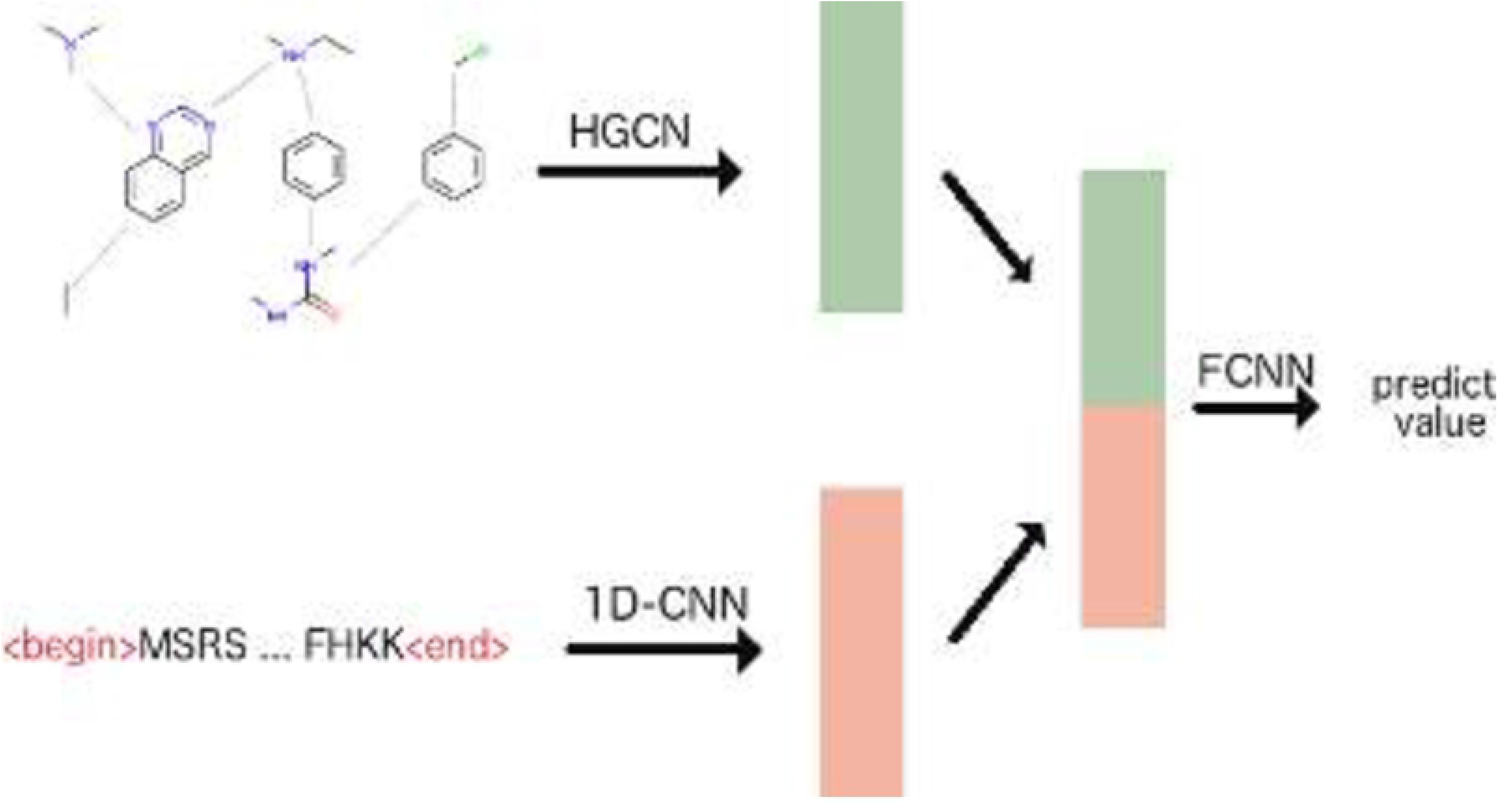
An overview of the CPI prediction model which encodes compounds using a HGCN and proteins using a 1D-CNN. FCNN denotes a fully-connected neural network, predicting the interaction based on a compound embedding and a protein embedding, which are generated from previously described components.

## Experiment and result

To evaluate our model and compare its performance to other deep learning based methods, we use four independent datasets. The compared methods include GCN [5], GraphDTA [10], TransformerCPI [3] and DeepDTA [11]. Among these methods, DeepDTA directly uses the SMILES format string as input and constructs a sequence-based model to encode compounds while the others employed different GCNs. DeepDTA and GraphDTA by default predict the binding affinity value between compounds and proteins. We modified their output layer from generating a float value into generating a binary value which allows us to directly compare the performance of these models.

The **Datasets** consist of *human* and *C*.*elegans* from [35], *bindingdb* created by [4] and *chembl27* downloaded from chEMBL database [40]. Except for the *chembl27* dataset, all datasets have been used to benchmark CPI prediction methods in previous studies [3–5, 7]. For the *human* and *C*.*elegans* datasets, Liu et al. [35] retrieved positive samples from two manually curated databases DrugBank 4.1 [41] and Matador [42], while they obtained negative samples using an in silico screening method. Gao et al. [4] created *bindingdb* dataset from BindingDB [43], a database of measured binding affinities, focusing primarily on the interactions of small molecules and proteins. The samples are labeled as positive if their IC50 is less than 100*nM* and negative if their IC50 is larger than 10, 000*nM*. For the *chembl27* dataset, we collected samples based on the binding activities of small molecules from the chEMBL database [44], only retaining the entries with a standard measure IC50. The thresholds to assign samples as positives or negatives are ≤ 100*nM* and ≥1000*nM* respectively. Table 1 shows the number of compounds, proteins, positive samples and negative samples for each dataset. With approximately 8000 samples the *human* and *C*.*elegans* datasets are substantially smaller than the other datasets, whereas they are more balanced in compound and protein quantity. Table 2 additionally offers an overview of the training, testing and validation dataset sizes. We evaluate the models on small datasets using k-fold cross validation, with *k* = 5. Meanwhile, the larger datasets are split into training, validation and testing subsets.

**Table 1.**
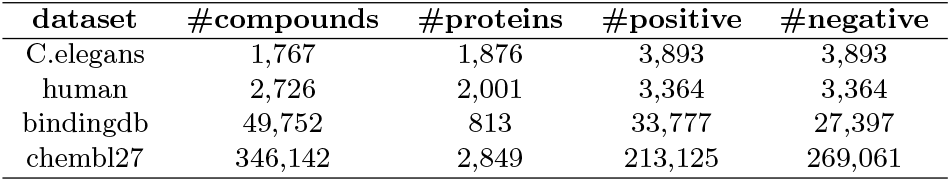
Summary of the experimental datasets.

| dataset | #compounds | #proteins | #positive | #negative |
| --- | --- | --- | --- | --- |
| C.elegans | 1,767 | 1,876 | 3,893 | 3,893 |
| human | 2,726 | 2,001 | 3,364 | 3,364 |
| bindingdb | 49,752 | 813 | 33,777 | 27,397 |
| chembl27 | 346,142 | 2,849 | 213,125 | 269,061 |

**Table 2.** Summary of the training and testing data. To small datasets, *human* and *C*.*elegans*, we evaluate the models’ performance using k-fold cross validation, with *k* = 5. To the other datasets, we split them into three sets: training, validation and testing.

|  | training |  | validation |  | testing |  |
| --- | --- | --- | --- | --- | --- | --- |
| dataset | #positive | #negative | #positive | #negative | #positive | #negative |
| C.elegans | - | - | - | - | - | - |
| human | - | - | - | - | - | - |
| bindingdb | 28,240 | 21,915 | 2,831 | 2776 | 2,802 | 2,706 |
| chembl27 | 171,022 | 214,728 | 21,076 | 27,142 | 21,027 | 27,191 |

Table 3 provides an insight to the testing data of all datasets and enumerates the number of samples in each case study, including new compound - known protein, known compound - new protein, new compound - new protein and known compound - known protein. A “known” compound/protein means the element has been seen in training data while a “new” compound/protein has the opposite meaning. Note that the known compound - known protein pair signifies an unseen pair built up from a compound and protein that are both in the training data, but never paired there. The last two columns of the table present the percentage of samples in the cases of new compound - new protein and of known compound - known protein. The testing data of *C*.*elegans* and *human* have very high rate of known compound - known protein samples, with 77.75% and 59.06%, respectively, while the rate of new compound - new protein samples are merely 0.8% and 3.59%, respectively. *bindingdb* and *chembl27* testing data have much lower rate of known-known samples, with less than 30%. They also contain a much higher rate of new samples, with 38.31% and 18.77%, respectively.

**Table 3.** A summary of CPIs in the testing data of two datasets *bindingdb* and *chembl27* in four scenarios: (a) new compound-known protein, (b) known compound-new protein, (c) new compound-new protein, and (d)known compound-known protein. The two last columns indicate percentage of (c) and (d) in testing data.

| dataset | (a) | (b) | (c) | (d) | %(c) | %(d) |
| --- | --- | --- | --- | --- | --- | --- |
| C.elegans | 174.8 (16.95) | 147.4 (8.65) | 12 (4.12) | 1168 (18.91) | 0.80 | 77.75 |
| human | 308.2 (4.55) | 155.8 (6.42) | 44.6 (5.23) | 733.8 (5.45) | 3.59 | 59.06 |
| bindingdb | 1682 | 397 | 2110 | 1319 | 38.31 | 23.95 |
| chembl27 | 18175 | 6,575 | 9051 | 14417 | 18.77 | 29.90 |

### Comparison in performance

For simplicity, all the customizable layers in the protein encoder were assigned the same kernel size. Validation data is used to tune the hyper-parameters, mainly for the kernel size in the customizable layers. The tuning process showed that *k* = 6 is sufficient for most experimental datasets. As a result, we choose *k* = 6 as the kernel size of our model in this experiment. We also trained our model with the same learning rate and weight decay, 1e−4, for all datasets. Table 4 shows MSE, F1 and AUC score of the prediction models on the experimental data. The performance of all methods is high on both *C. elegans* and *human* data, with *F*1 scores higher than 0.90 for both datasets. Meanwhile, performance on *bindingdb* and *chembl27* is much lower for all models. For example, the GCN method achieves approximately 0.700 in terms of F1 score on these datasets. These results can be explained by the degree of overlap in proteins and/or compounds between the training and the testing data, shown in Table 3. Our method works only better than the GCN method on *C*.*elegans* data but it shows a similar performance to DeepDTA and better than the others on *human* data. Furthermore, it outperforms GraphDTA and TransformerCPI on two large datasets, *bindingdb* and *chembl27*. In the point of average ranking, our method outperforms the other methods in predicting CPIs.

**Table 4.** MSE, F1 and AUC scores of CPI prediction models on the experimental data.

| MSE |  |  |  |  |  |
| --- | --- | --- | --- | --- | --- |
|  | GCN | GraphDTA | TransformerCPI | DeepDTA | Ours |
| C.elegans | 0.209 | 0.140 | 0.169 | 0.093 | 0.156 |
| human | 0.253 | 0.197 | 0.281 | 0.171 | 0.180 |
| bindingdb | 0.574 | 0.431 | 0.335 | 0.331 | 0.272 |
| chembl27 | 0.525 | 0.518 | 0.446 | 0.470 | 0.415 |
| average rank | 4.75 | 3.25 | 3.5 | 1.75 | 1.75 |
| F1 |  |  |  |  |  |
|  | GCN | GraphDTA | TransformerCPI | DeepDTA | Ours |
| C.elegans | 0.932 | 0.957 | 0.946 | 0.973 | 0.950 |
| human | 0.915 | 0.934 | 0.906 | 0.944 | 0.947 |
| bindingdb | 0.706 | 0.810 | 0.858 | 0.875 | 0.895 |
| chembl27 | 0.679 | 0.690 | 0.772 | 0.751 | 0.797 |
| average rank | 4.75 | 3.25 | 3.5 | 2 | 1.5 |
| AUC |  |  |  |  |  |
|  | GCN | GraphDTA | TransformerCPI | DeepDTA | Ours |
| C.elegans | 0.976 | 0.989 | 0.986 | 0.994 | 0.985 |
| human | 0.969 | 0.978 | 0.960 | 0.984 | 0.983 |
| bindingdb | 0.881 | 0.886 | 0.933 | 0.951 | 0.959 |
| chembl27 | 0.814 | 0.820 | 0.883 | 0.865 | 0.896 |
| average rank | 4.5 | 3 | 3 | 2.5 | 2 |

To allow a more detailed evaluation, we applied the trained models on the testing data, specifically on four subsets corresponding to four different scenarios, as mentioned in Table 3. Table 5 presents the AUC scores of the methods in each scenario for each dataset. In general, DeepDTA outperforms the other methods in predicting CPIs with known compound and known protein. In this scenario, our method outperforms the remaining ones, including GCN,TransformerCPI, and GraphDTA. For new compound - known protein samples, our model achieves a performance similar to DeepDTA and GraphDTA, obtaining 2.5 in average ranking. Furthermore, it outperforms the others in terms of average ranking in the other scenarios, known compound - new protein and new compound - new protein.

**Table 5.** AUC score of the methods in four scenarios for each dataset.

| new compound - known protein |  |  |  |  |  |
| --- | --- | --- | --- | --- | --- |
|  | GCN | GraphDTA | TransformerCPI | DeepDTA | Ours |
| C.elegans | 0.907 | 0.972 | 0.969 | 0.971 | 0.948 |
| human | 0.923 | 0.969 | 0.964 | 0.947 | 0.962 |
| bindingdb | 0.971 | 0.960 | 0.957 | 0.982 | 0.981 |
| chembl27 | 0.867 | 0.900 | 0.939 | 0.921 | 0.942 |
| average rank | 4.5 | 2.5 | 3 | 2.5 | 2.5 |
| known compound - new protein |  |  |  |  |  |
|  | GCN | GraphDTA | TransformerCPI | DeepDTA | Ours |
| C.elegans | 0.978 | 0.990 | 0.981 | 0.997 | 0.993 |
| human | 0.975 | 0.961 | 0.915 | 0.996 | 0.987 |
| bindingdb | 0.758 | 0.585 | 0.778 | 0.831 | 0.850 |
| chembl27 | 0.771 | 0.582 | 0.737 | 0.746 | 0.776 |
| average rank | 3.5 | 4.25 | 4 | 1.75 | 1.5 |
| new compound - new protein |  |  |  |  |  |
|  | GCN | GraphDTA | TransformerCPI | DeepDTA | Ours |
| C.elegans | 0.334 | 0.851 | 0.700 | 0.929 | 0.772 |
| human | 0.635 | 0.776 | 0.699 | 0.783 | 0.756 |
| bindingdb | 0.840 | 0.792 | 0.897 | 0.895 | 0.915 |
| chembl27 | 0.708 | 0.625 | 0.697 | 0.692 | 0.740 |
| average rank | 4 | 3.5 | 3.25 | 2.25 | 2 |
| known compound - known protein |  |  |  |  |  |
|  | GCN | GraphDTA | TransformerCPI | DeepDTA | Ours |
| C.elegans | 0.983 | 0.992 | 0.988 | 0.997 | 0.990 |
| human | 0.985 | 0.985 | 0.969 | 0.995 | 0.992 |
| bindingdb | 0.974 | 0.955 | 0.944 | 0.993 | 0.986 |
| chembl27 | 0.829 | 0.884 | 0.938 | 0.924 | 0.948 |
| average rank | 4.125 | 3.375 | 4 | 1.5 | 2 |

### Effect of the kernel size

In this experiment, we examine how the kernel size in 1D-CNN layers and max pooling layers affects our model performance. We changed the kernel size in the customizable layers to 6, 8, 10, 12 and trained different versions of our model. Figure 5 shows AUC scores of these trained predictors on testing data for two datasets: *human* and *bindingdb*. For the *human* dataset, the AUC score slightly changes as a function of the kernel size, reaching the highest value at the kernel size *k* = 10. The AUC value of the model trained with the kernel size *k* = 6 is also promising, which is lower than the highest value merely 0.1%. For the *bindingdb* dataset, the performance generally decreases when the kernel size increases from *k* = 6 to *k* = 12, with an AUC difference of 3%. This indicates that kernel size can have an effect on the performance of the model. The kernel size defines the number of adjacent amino acids in proteins that would be considered in one patch. As the ideal split can be different among proteins, the ideal kernel size is also different among datasets. With the result we obtained from four experimental datasets, a kernel size of 6 is a good option to use.

**Fig 5.**
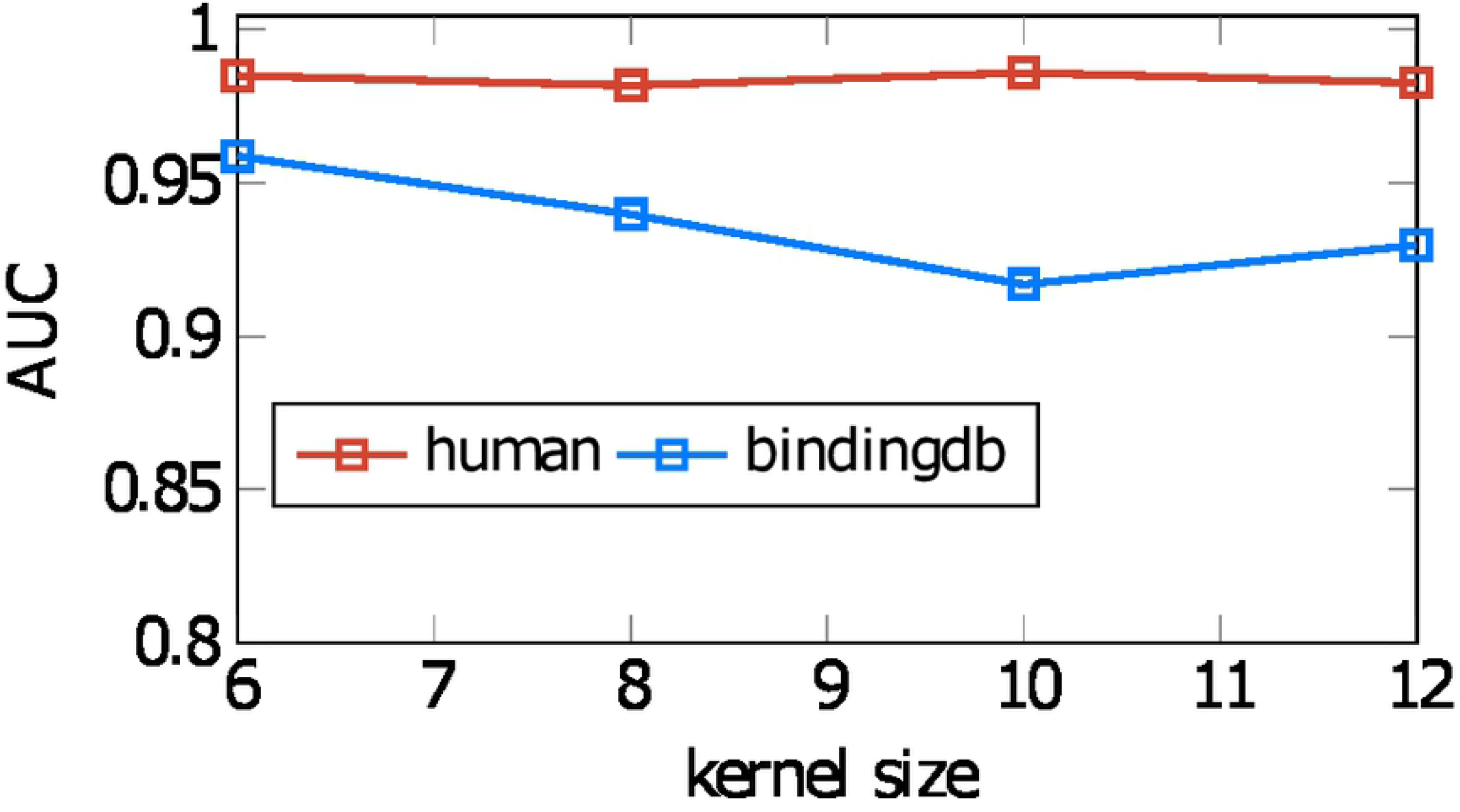
Performance of our model on two datasets, *human* and *bindingdb*, when it is trained with different 1D-CNN kernel sizes.

To get an insight on how the kernel size impacts our model, we also examined the performance of our model trained on different kernel size in the four different scenarios. Figure 6 and 7 show the result of the examination on *human* and *bindingdb* data, respectively. In the scenarios of new compound - known protein and known compound - known protein, our results indicate slight differences in AUC score of our model when it is trained with the kernel size of 6, 8, 10 and 12. In other words, changing the kernel size does not affect much on the prediction of CPIs with known proteins although the proteins are paired with new compounds. This is reasonable as the kernel size is merely related to the protein encoder. A large change in performance when modifying the kernel size can be observed for the scenario of new compound - new protein. The kernel size of 6 results in the highest AUC score, producing a model that is able to recognize completely new CPIs better than models of the other kernel values.

**Fig 6.**
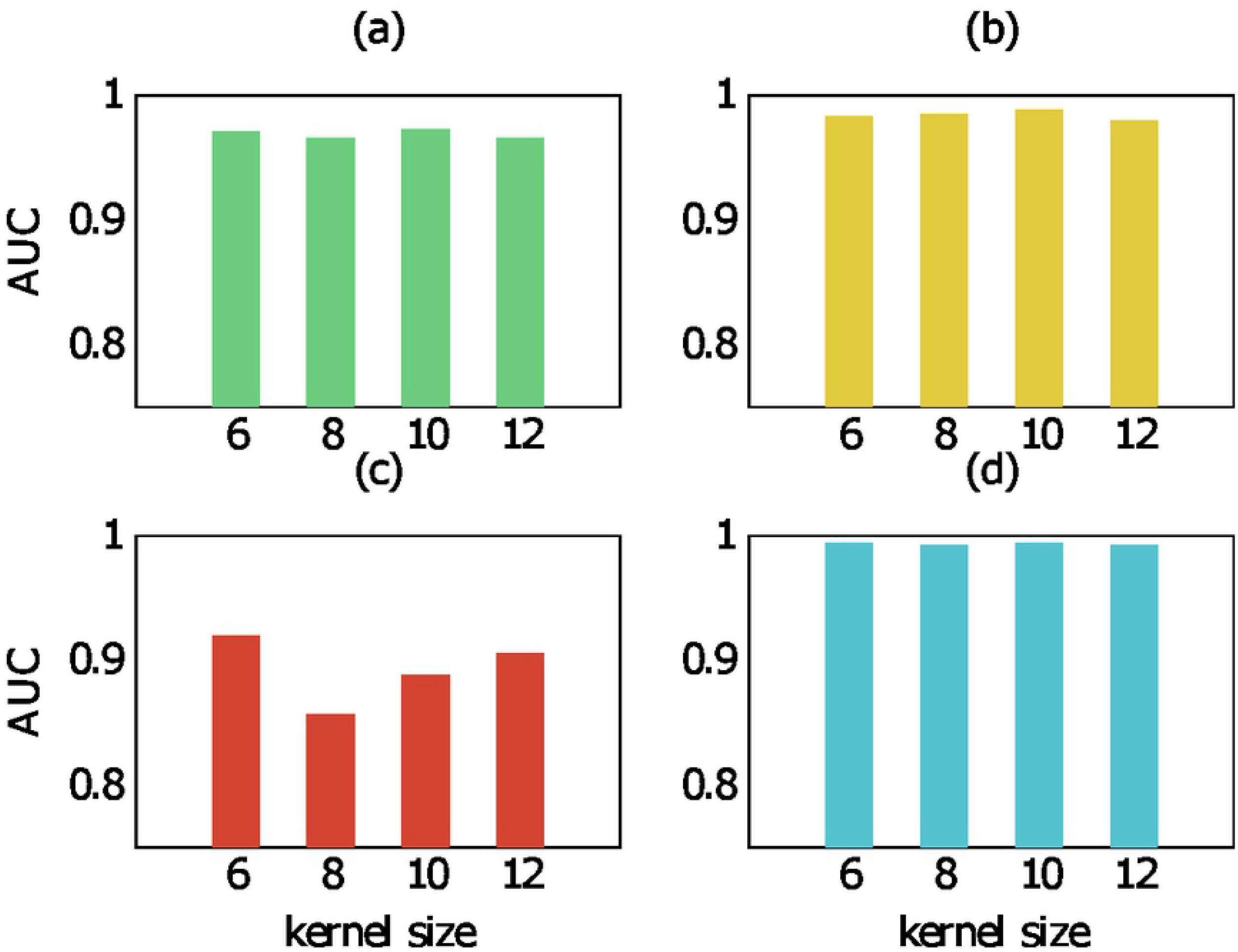
AUC score of our method trained with different kernel size on human data in four scenarios:(a)new compound-known protein, (b)known compound-new protein, (c)new compound-new protein, (d) known compound-known protein.

**Fig 7.**
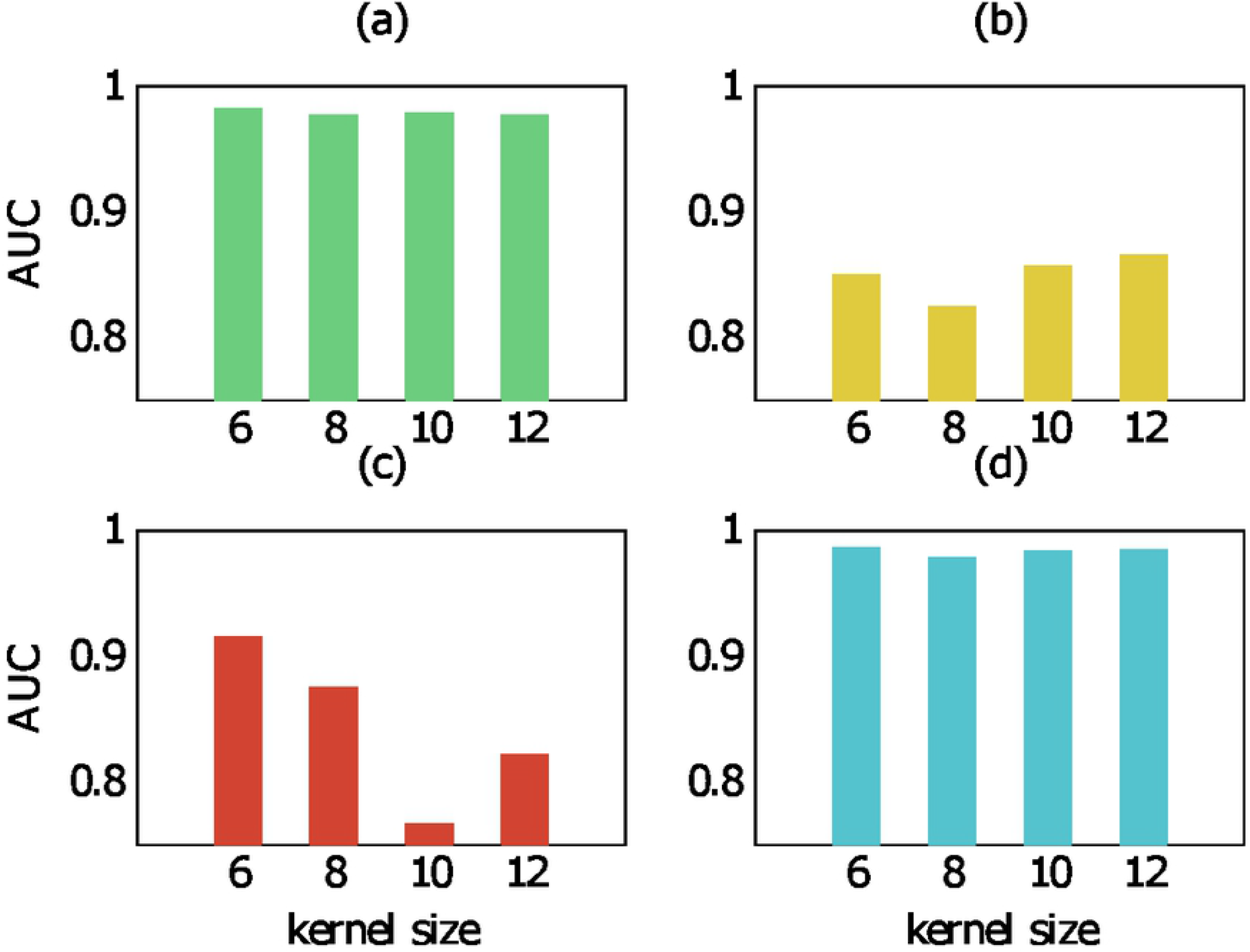
AUC score of our method trained with different kernel size on bindingdb data in four scenarios:(a)new compound-known protein, (b)known compound-new protein, (c)new compound-new protein, (d) known compound-known protein.

## Explanation with Grad-CAM

Grad-CAM [36] can be used to produce a visual explanation of the decisions made by CNN-based models, without modifying the base models or requiring re-training. Grad-CAM is applicable to any CNN-based architecture, including those for image captioning and visual question answering. Grad-CAM operates by flowing gradient information back to the last convolutional layer, where importance values are assigned to each neuron for a particular decision of interest. These importance values are used to compute a class-discriminative localization map. To obtain the full explanation, we can interpolate the map and project the result onto the original input. For 2D-CNN-based models, the class-discriminative localization map Grad-CAM 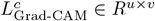 of width *u* and height *v* for any class *c* is computed as follows:

- Neuron importance weights 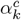 are computed using the gradient of the score for class *c, y*^*c*^, with respect to the feature map activations *A*^*k*^ in the last convolutional layer, *i*.*e*.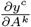.

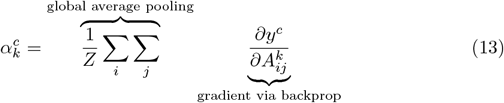
- The neural importance weight 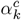 captures the “importance” of feature map *k* for the target class *c*. These are then combined with forward activation maps, followed by a RELU function to produce,

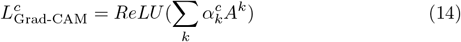 The ReLU is applied to the linear combination of maps in order to focus only on the features that have a positive influence on the class of interest.

For the 1D-CNN-based model, the equation 13 is replaced by:

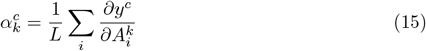

where *L, A*^*k*^ denote the length and the whole dimension *k*-th of the feature map, respectively.

Figure 8 is an example of a class-discriminative localization map 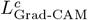 that Grad-CAM explains for a CPI prediction generated by our model. This example represents the interaction between Protein Kinase R-like Endoplasmic Reticulum Kinase (PERK) and the small molecule 3Z1 (PDB ID 4X7J). PERK is one of three sensors of misfolded proteins that are known to mediate the unfolded protein response (UPR) through complementary pathways [45]. The two heat maps correspond to two different models trained with kernel size *k* = 6 and *k* = 12, respectively. The heat maps reveal the residues predicted to play the most important role in the interaction. With a smaller kernel size, each signal at the last 1D-CNN layer is associated with a smaller region in the input. Therefore, we obtain a more detailed explanation with a smaller kernel size, focusing on shorter regions, compared to larger kernel sizes. Figure 9 shows the 3D visualization of the same CPI in which (A) is the structure obtained from PDB, while (B) and (C) highlight the importance of the residues according to the prediction models trained with kernel size *k* = 6 and *k* = 12, respectively. The prediction models have concentrated on several binding site residues that are within 5Å from the ligand when making their decision. However, other residues were also taken into account that are far from the ligand. When comparing the two prediction models, the model with kernel size *k* = 6 is focused more on the residues surrounding the ligand in a radius of 5Å than the model with the kernel size *k* = 12.

**Fig 8.**
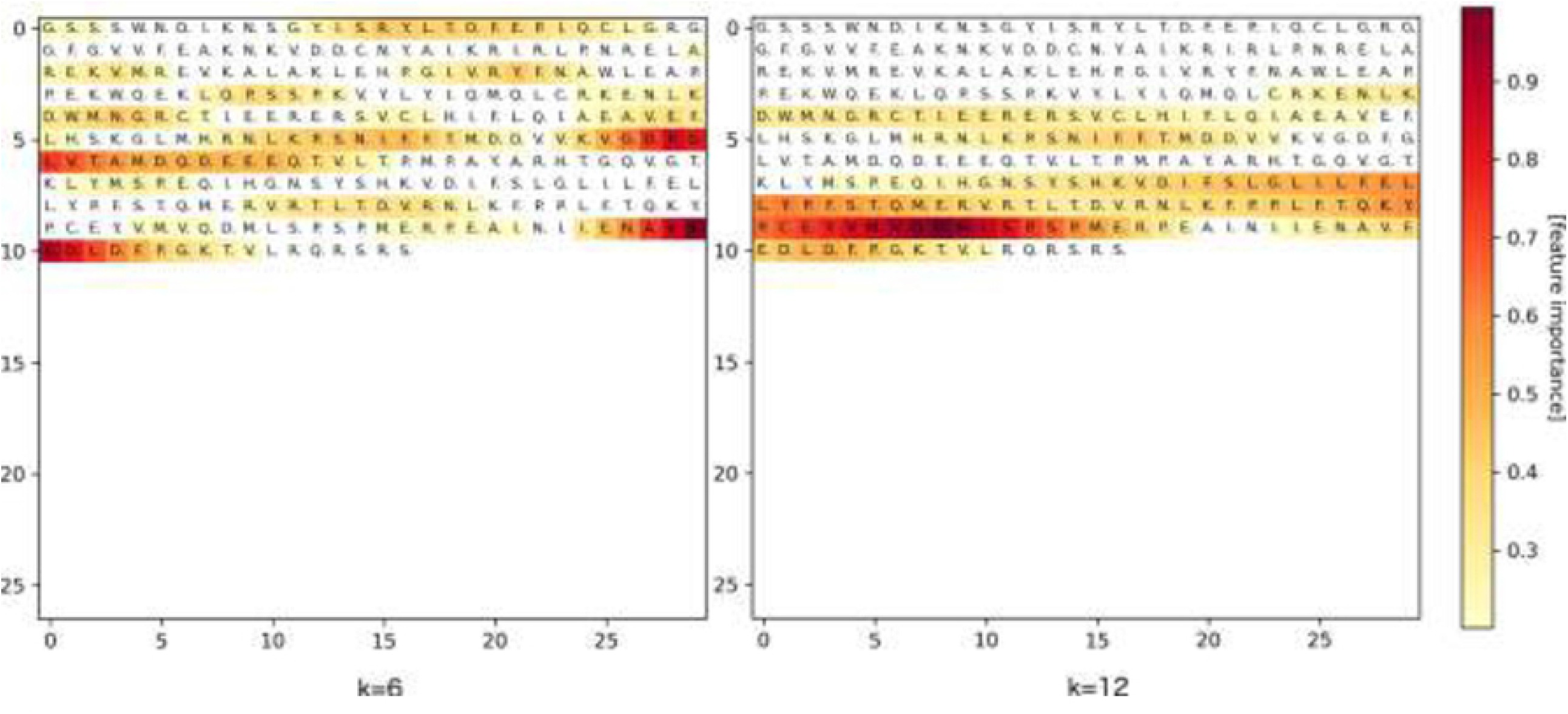
An example of a visual explanation generated by Grad-CAM for a CPI prediction made by our model. The CPI occurs between the protein PERK and the ligand 3Z1 (PDB ID 4X7J). The left plot corresponds to the model trained with kernel size *k* = 6 and the right plot to a model with kernel size *k* = 12. The darker the residues are, the more importance these amino acids contribute to the prediction.

**Fig 9.**
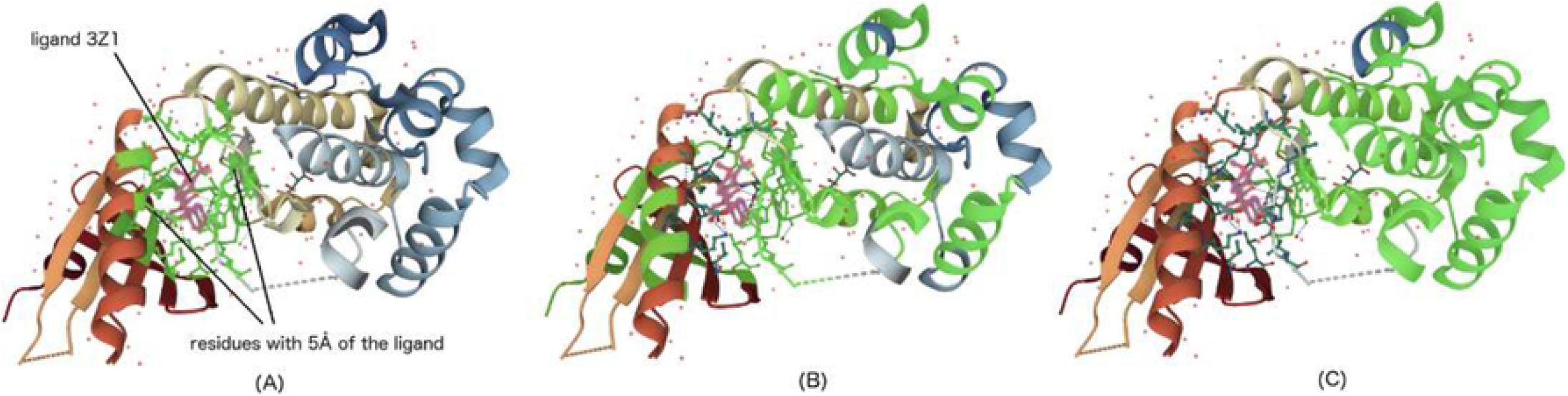
3D visualization of CPI between PERK and the ligand 3Z1 (PDB ID 4X7J): (A) structure retrieved from PDB; (B) explanation from the model with kernel size *k* = 6; and (C) explanation from the model with kernel size *k* = 12. The central structure in pink color is the ligand 3Z1. The green part in (A) denote the residues located within 5Å of the ligand, whereas the green regions in (B) and (C) correspond to the residues that contribute most (≥0.2) to the CPI prediction. The contribution values are taken from the corresponding heat maps in figure 8 with cut off value 0.2.

## Conclusion

We proposed a new hierarchical graph convolutional network based technique which is able to produce three levels of embedding: node, subgraph and entire graph. We used this GCN model as a compound encoder, combining it with a 1D-CNN based model in order to build a complete CPI prediction framework. To generate subgraphs in compounds, we developed a simple procedure based on aromatic links and connected components. The experimental results demonstrate that our framework outperforms existing GCN based models on different CPI datasets. Compared to DeepDTA, a highly performing sequence-based CPI prediction method, our model is competitive, showing improved performance when large training data is available. In addition, the visual explanation provided by Grad-CAM shows the relevance of the involved residues for the proposed model. This offers a promising extension towards the prediction of binding sites between compounds and proteins.

We employed different types of neural network layers to build the CPI prediction model, recruiting basic information from compounds and proteins as input. The embedding of compounds and proteins are learnt merely from CPI data during the training process. However, there is a large number of small molecules and protein sequences available in public databases that can be exploited to learn compound and protein embeddings through unsupervised approaches. Pre-trained embedding has the potential to make learning significantly easier and cheaper. In addition, attention mechanisms have proven their efficiency in several deep learning models. Integrating a relevant attention mechanism into our CPI prediction framework may further improve its performance. We leave this as an open avenue for exploration in future work. The visual explanation we provide on the protein sequences suggests possible binding sites which offers opportunities to generate new biological insights in protein function and opens applications in the context of drug discovery and re-purposing. As the incorporation of compound substructure information yields good CPI prediction performances, we argue that substructure analysis can be a valuable step in computational drug design.

